# The Arabian camel, *Camelus dromedarius* interferon epsilon: functional expression, *in vitro* refolding, purification and cytotoxicity on breast cancer cell lines MDA-MB-231 and MCF-7

**DOI:** 10.1101/568329

**Authors:** Manal Abdel-Fattah, Hesham Saeed, Lamiaa El-Shennawy, Manal Shalaby, Amira M. Embaby, Farid Ataya, Hoda E.Mahmoud, Ahmed Hussein

## Abstract

The current study highlights for the first time cloning, overexpression, purification, and assessing the cytotxcity of the novel interferon epsilon (IFNε), from the Arabian camel *Camelus dromedarius*, against two human breast cancer cell lines MDA-MB-231 and MCF-7. Full-length cDNA encoding interferon epsilon (IFNε) was isolated and cloned from the liver of the Arabian camel, *C. dromedarius* using reverse transcription-polymerase chain reaction. The sequence analysis of the camel IFNε cDNA showed a 582-bp open reading frame encoding a protein of 193 amino acids with an estimated molecular weight of 22.953 kDa. A BLAST search analysis revealed that the *C. dromedarius* IFNε shared high sequence identity with the IFN genes of other species, such as *Camelus ferus*, *Vicugna pacos*, and *Homo sapiens*. Expression of the camel IFNε cDNA in *Escherichia coli* gave a fusion protein band of 22.73 kDa after induction with either isopropyl β-D-1-thiogalactopyranoside or lactose for 5 h. Recombinant IFNε protein was overexpressed in the form of inclusion bodies that were easily solubilized and refolded using SDS and KCl. The solubilized inclusion bodies were purified to apparent homogeneity using nickel affinity chromatography. We examined the effect of IFNε on two breast cancer cell lines MDA-MB-231 and MCF-7. In both cell lines, IFNε inhibited cell survival in a dose dependent manner as observed by MTT assay, morphological changes and apoptosis assay. Caspase-3 expression level was found to be increased in MDA-MB-231 treated cells as compared to untreated cells.

## Introduction

The term interferon (IFN) was first described by Alick Isaacs and Jean Lindemann in 1957 at the National Institute for Medical Research in London. They described an antiviral agent produced by virally infected chick cells, and they called it interferon (IFN), a substance that interferes with influenza and vaccinia virus replication [1]. Since then, the efforts to discover, characterize, and develop new interferons as major therapeutic proteins have continued for over 60 years [2]. IFNs are members of a large cytokine family of evolutionarily conserved pleiotropic regulators of cellular functions; they are relatively low-molecular weight signaling proteins (20–25 kDa) usually glycosylated and produced by a variety of cells, such as epithelia, endothelia, stroma, and cells of the immune system [3–5]. The expression of IFNs is induced by a variety of different stimuli associated with viral infections, bacteria, parasites, inflammation, and tumorigenesis [6]. IFNs, therefore, induce a diverse range of biological functions and responses, including cell proliferation and differentiation, inflammation, chemotaxis, immune cell (natural killer cells and macrophages) activation, and apoptosis [7, 8]. The key to understanding these regulatory proteins lies in the recognition of their pleiotropism, overlapping activities, functional redundancies, and side effects [3]. Based on the type of receptors they interact with for signal transduction, IFNs are classified into three major types namely, type I, II, and III, which have different gene and protein structures and biological activities [9]. The mammalian type I IFNs represents a large family of related proteins, mainly virus-inducible, divided into eight subfamilies named α, β, ω, δ, ε, ν and κ [10, 11]. Besides the autocrine activation of antiviral responses, type I IFNs function systematically to induce an antiviral state in the surrounding and distal cells [12, 13]. In combination with chemo and radiation therapies, interferon therapy is used as a treatment of some malignant diseases, such as hairy cell leukemia, chronic myeloid leukemia, nodular lymphoma, and cutaneous T-cell lymphoma [14]. The recombinant IFN-α2b can be used for the treatment of patients with recurrent melanomas [15]. Hepatitis B, hepatitis C, and HIV are treated with IFN-α often in combination with other antiviral drugs [16, 17].

One of the most recently discovered interferon is the interferon epsilon (IFNε). Signal transduction by IFNε is mediated through binding to the interferon α/β receptor (IFNAR), despite its low sequence homology with α- and β-type interferons. Although binding to the same heterodimeric receptor pair, they evoke a broad range of cellular activities, affecting the expression of numerous genes and resulting in profound cellular changes [12, 18–20]. The expression of IFNε is neither induced by a pattern recognition receptor pathway nor by an exposure to viral infection [21]. Unlike other type-I IFNs, IFNε is constitutively expressed in the lung, brain, small intestine, and reproductive tissue; thus, it is thought to play a role in reproductive function, in either viral protection or early placental development in placental mammals [18, 19]. IFNε has high amino acid sequence homology with other type-I interferons, of which IFN-β is the closest paralog, and they share 38% identical residues. A common structural feature of IFNε is the lack of a disulfide linkage and the presence of two glycosylation sites represented by asparagine 74 and 83. Many IFNs genes have been cloned and characterized from a variety of species such as human, pig, mouse, dog, cat, cattle, chicken, turkey, goose, zebra fish, and Atlantic salmon [22–25]. However, the information about the IFNε from the Arabian one-humped camel, *Camelus dromedarius*, has not been reported yet. This domesticated camel is one of the most important animals in the Arabian Peninsula, having high cultural and economic value. In Saudi Arabia, it comprises 16% of the animal biomass and is considered as the main source of meat [26, 27]. The aim of the present study was the isolation of full-length *C. dromedarius IFNε* gene, followed by its expression in *Escherichia coli*, *in vitro* refolding of the recombinant protein, purification, and characterization of the purified IFNε protein. Cytotoxicity and apoptosis assays were then performed to define the effect of the purified recombinant IFNε protein on human cancer cell lines. The results of this study contribute towards the importance of discovering and characterizing IFNε from this unique Arabian camel, and propose its potential use for the treatment of cancer.

## Materials and methods

### Chemicals and reagents

All chemicals and reagents were of molecular biology, analytical, or chromatographic grade. Water was de-ionized and milli-Q-grade.

### Tissue collection and RNA isolation

Liver tissues (2 g) from adult male one-humped Arabian camel, *C. dromedarius*, were collected from a slaughter house located in the north of Riyadh City, Kingdom of Saudi Arabia. The animals were sacrificed under the observation of a skilled veterinarian, and the liver samples were taken and immediately submerged in 5 mL of RNA later^®^ solution (Ambion, Courtabeuf, France) to preserve the integrity of RNA. The samples were kept at 4 °C overnight and thereafter stored at −80 °C until used for RNA isolation. The liver samples were removed from −80 °C and left at room temperature until thawed completely. Fifty milligrams were homogenized in 0.5 mL RLT lysis solution supplemented with 1% 2-mercaptoethanol using a rotor-stator homogenizer (MEDIC TOOLS, Switzerland). Total RNA was isolated and purified using the RNeasy Mini Kit (Qiagen, Germany), with a DNase digestion step following the manufacturer’s protocol. The elution step was performed using 50 μL nuclease free water. The concentration, purity, and integrity of the isolated purified total RNA were determined using the Agilent 2100 Bioanalyzer System and Agilent total RNA analysis kit, according to the manufacturer’s protocols (Agilent Technologies, Waldbronn, Germany).

### First strand cDNA synthesis and amplification of camel IFNε gene

Total RNA, isolated previously from adult male one-humped Arabian camel, *C. dromedaries*, was used in the current study as a source for camel *IFNε* gene. Two micrograms of total RNA were reverse transcribed into the first strand cDNA using the ImProm-II Reverse Transcription System (A3800, Promega, Madison, USA) according to the manufacturer’s protocol and used as a template for the amplification of the full-length camel IFNε cDNA. A polymerase chain reaction (PCR) was conducted in a final volume of 50 μL, containing 25 μL 2X high-fidelity master mix (GE Healthcare, USA), 3 μL (30 pmol) of each *IFNε* gene forward primer that contains an *Eco*RI restriction site (5’-<u>GAATTC</u> ATGATTAACAAGCCTTTCTT-3′) and a reverse primer that contains a *Hind*III restriction site (5′-<u>AAGCTT</u>AGGATCCATTCCTTGTTTGC-3′), and 5 μL cDNA. The PCR amplification was performed using the following reaction conditions: 1 cycle at 95 °C for 5 min, followed by 30 cycles at 95 °C for 30 s, 55 °C for 30 s, and 72 °C for 1 min. A final extension step was carried out at 72 °C for 5 min. The PCR products were resolved in a 1.5% agarose gel stained with 0.5 μg/mL ethidium bromide.

### Cloning and sequencing of the full-length camel IFNε cDNA

The PCR product was first cloned into the pGEM^®^-T Easy vector (Promega Co. Cat #A1360) to facilitate the sequencing process and subcloning into the pET28a (+) expression vector. The ligation reaction was carried out in a clean sterile 1.5-mL eppendorf tube containing 4 μL of the PCR product, 1 μL (50 ng) of pGEM^®^-T-Easy vector (Promega, USA), 1 μL of 10X ligase buffer, and 1 U of ligase enzyme. The final volume of the reaction was adjusted to 10 μL by the addition of nuclease free water. The reaction tubes were kept at 16 °C overnight, after which 5 μL was used to transform the *E. coli* JM109 competent cells, according to Sambrook et al. (1989) [28]. Screening was carried out on the selective LB/IPTG/X-gal/Ampicillin/agar plates. The recombinant plasmids were prepared from some positive clones using the PureYield Plasmid Miniprep System (Cat #A1222, Promega, Madison, USA). The sequencing of the cloned insert was carried out according to Sanger et al. 1977 [29] using the T7 (5′-TAATACGACTCACTATAGGG-3′) and SP6 (5′- TATTTAGGTGACACTATAG-3′) sequencing primers. The sequence analysis was carried out using the DNAStar, BioEdit, and Clustal W programs.

### Phylogenetic tree and structure modeling analysis

A phylogenetic tree analysis was constructed according to Dereeper et al. [30], using the Phylogeny.fr software (http://www.Phylogeny.fr). The nucleotide sequences for the Arabian camel IFNε cDNA was analyzed using the basic local alignment search tool (BLAST) programs BLASTn, BLASTp (http://www.ncbi.nlm.nih.gov), and a multiple sequence alignment was carried out using the ClustalW, BioEdit, DNAStar, and Jalview programs. The protein sequence was obtained by translating the cDNA nucleotides sequence by using a translation tool at the ExPasy server (http://web.expasy.org/translate/). The protein sequence was submitted to the Swiss-Model server for structure prediction, and the structural data were analyzed by the PDB viewer program. Finally, the predicted 3D structure models were built based on the multiple threading alignments by using the local threading meta-server (LOMET) and iterative TASSER assembly simulation [31, 32].

### Sub-cloning into pET-28a (+) vector

The IFNε cDNA insert cloned into the pGEM-T-Easy plasmid was released using the *Eco*RI and *Hind*III restriction enzymes (2 units each) according to Sambrook et al. (1989) [28]. The released insert was purified from the agarose gel using the QIAquick Gel Extraction Kit (Cat. # 28704, QIAGEN) and sub-cloned into the pET-28a (+) expression vector. The plasmid pET-28a (+) (Novagen) carries an N-terminal His-Tag/thrombin/T7 configuration, and the expression of the cloned gene is under the control of a T7 promoter. A 2-μg aliquot of plasmid pET-28a (+) was digested with 2 units of *Eco*RI and *Hind*III at 37 °C overnight, after which the digestion reaction was terminated by heating the tubes at 65 °C for 15 min. The linearized plasmid was treated with 2 units of shrimp alkaline phosphatase (Promega, Madison, USA) at 37 °C for 30 min. Finally, the reaction was terminated by incubation at 70 °C for 10 min. The ligation reaction was carried out in a tube containing 2 μL (50 ng) of pET28a (+), 2 μL (100 ng) of IFNε cDNA insert, 1 μL 10X ligase buffer, and 1 μL (2 units) of ligase enzyme. The final volume was adjusted to 10 μL by the addition of nuclease free water, and the tube was incubated at 16 °C overnight. Subsequently, 5 μL of the ligation reaction was used to transform *E. coli* BL21(DE3) pLysS (Cat. # P9801, Promega, USA) competent cells, according to Sambrook et al. (1989) [28]. The recombinant *E. coli* BL21(DE3) pLysS harboring the pET-28a (+) vector was screened on the selective LB/IPTG/X-gal/Kanamycin/agar plates and by using the colony PCR strategy utilizing the IFNε gene-specific primers. The recombinant plasmids were isolated from the positive clones using the Pure Yield Plasmid Miniprep System (A1222, Promega, USA), and some potential positive plasmids containing the cDNA insert were digested with *Eco*RI and *Hind*III to confirm the presence of the IFNε cDNA insert.

### Expression of camel IFNε cDNA in *E. coli* BL21(DE3) pLysS

The transformed *E. coli* BL21(DE3) pLysS harbouring the recombinant plasmid were cultured in 1 L of Luria broth medium supplemented with 34 μg/mL kanamycin and incubated at 37 °C for 4 h at 250 rpm. When the optical density at 600 nm reached 0.6, isopropyl-β-D-1-thiogalactopyranoside (IPTG) was added to the culture at a concentration of 1 mM. The culture flask was incubated at 37 °C with shaking at 250 rpm for 5 h, after which the bacterial cells were harvested by centrifugation at 8000 rpm for 20 min at 4 °C. The bacterial pellets were re-suspended in 10 mL of 0.1 M potassium phosphate buffer, pH 7.5, containing 50% glycerol. The bacterial cell suspension was then sonicated on an ice-bath using 4x 30-s pulses, and the cell debris were removed by centrifugation at 10,000 rpm for 10 min at 4 °C, after which the supernatant and pellets were collected in separate eppendorf tubes. The pellets were re-suspended in 5 mL of 0.1 M potassium phosphate buffer, pH 7.5, containing 50% glycerol and both supernatant and pellets were used for further analysis. The gene expression was also analyzed using lactose as an inducer at a concentration of 2 g/L in the fermentation medium.

### Protein determination

Protein concentration was determined according to Bradford (1976) [33], using 0.5 mg/mL of bovine serum albumin (BSA) as a standard.

### Sodium dodecyl sulfate gel electrophoresis (SDS-PAGE)

The expression of the camel recombinant *IFNε* gene in *E. coli* was checked by performed a 12% SDS-PAGE according to Laemmli, 1970 [34]. After electrophoresis, the gel was stained with Coomassie Brilliant Blue R-250 followed by de-staining in a solution of 10% (v/v) methanol and 10% (v/v) acetic acid.

### Refolding of *C. dromedarius* recombinant IFNε protein inclusion bodies

The transformed *E. coli* BL21(DE3) pLysS cells containing over-expressed camel IFNε protein was disrupted by sonication, and the inclusion bodies were recovered by centrifugation at 10,000 rpm for 30 min at 4 °C. The pellets were washed three times with 20 mM Tris-HCl, pH 8.0, and after the final wash, the pellets were resuspended in denaturation buffer containing 50 mM M Tris-HCl (pH 8.0), 0.3 M NaCl, and 2% SDS with continuous stirring on an ice-bath until the solution becomes clear. The protein solution was then kept at 4 °C overnight, followed by centrifugation for 10 min at 10,000 rpm and 4 °C to precipitate the excess SDS. Subsequently, KCl was added to the supernatant at a final concentration of 400 mM, and the solution was kept at 4 °C overnight. Thereafter, centrifugation was carried out for 10 min at 10,000 rpm and 4 °C, and the clear solution was dialyzed overnight against 50 mM potassium phosphate buffer (pH 7.5) and applied to a nickel affinity column [35, 36].

### Affinity purification of *C. dromedarius* recombinant IFNε

The recombinant IFNε protein was purified using a single-step High-Select High Flow (HF) nickel affinity chromatographic gel (Sigma-Aldrich, Cat. # H0537). The nickel affinity column (1.0 cm × 1.0 cm) was packed with the affinity matrix and washed thoroughly with 30 mL of de-ionized water, followed by equilibration with the 5-bed volumes of 50 mM potassium phosphate buffer (pH 7.5) containing 20 mM imidazole. A solution of solubilized inclusion bodies (5 mL) was loaded onto the column, and the column was washed with 5-bed volumes of 50 mM potassium phosphate buffer (pH 7.5) containing 20 mM imidazole. The recombinant IFNε protein was eluted with 50 mM potassium phosphate buffer (pH 7.5) containing 500 mM imidazole. The collected fractions were measured at 280 nm, and the fractions presented in the second peak were pooled together and dialyzed overnight against 50 mM potassium phosphate buffer (pH 7.5). The purity of the dialyzed recombinant IFNε protein was checked by performing 12% SDS-PAGE.

### Electron microscopy analysis

The recombinant *C. dromedarius* IFNε inclusion bodies were fixed in a solution of formaldehyde and glutaraldehyde (4:1) and observed and analyzed by transmission electron microscopy (TEM; JEOL-JSM 1400 plus) and scanning electron microscopy (SEM; JEOL-JSM 5300).

### Cytotoxicity of *C. dromedarius* recombinant IFN epsilon on breast cancer cell lines

Human breast cancer cell lines, MDA-MB-231 and MCF-7, were obtained from the lab of Professor Stig Linder, Karolinska Institute, Sweden. Cells were cultured in Dulbecco’s Modified Eagle’s Medium supplemented with 10% Fetal Bovine Serum (Sigma), 100 U/mL penicillin, and 100 mg/mL streptomycin. Cells were maintained in 5% CO_2_ at 37 °C.

### MTT assay

MDA-MB-231 and MCF-7 cells were seeded in 96 well plates (15,000 and 10,000 cells/ well, respectively). After 24 h, cells were treated with different concentrations of recombinant interferon epsilon and the control cells received untreated medium in the same buffer. Cells were washed twice with PBS after 48 h of incubation, and 3-(4,5-Dimethyl-2-thiazolyl)-2,5-diphenyl-2H-tetrazolium bromide (MTT) reagent (10 μL of 5 mg/mL) (Serva) in 100 μL serum free medium was added to each well. After 3-4 h of incubation at 37 °C, the medium was discarded, and cells were incubated with 100 μL of DMSO. Plates were shaken, then absorbance was measured at 490 nm [37].

### Apoptosis assay

Apoptosis was analyzed using Annexin V-FITC apoptosis detection kit (Miltenyi Biotec). MDA-MB-231 and MCF-7 were incubated with recombinant IFNε for 48 h. The floating cells were detached from the plate surface and attaching cells were harvested by trypsinization and pelleted by centrifugation. The cell pellets were resuspended in binding buffer and incubated with fluorescein isothiocyanate (FITC)-labeled Annexin V for 15 min in the dark at room temperature. Cells were washed and resuspended in binding buffer, then and Propidium Iodide was added. The stained cells were analyzed in Flow Cytometry Service core at Center of Excellence for Research in Regenerative Medicine and its Applications using BD FACSCalibur flow cytometer (BD Biosciences).

### Caspase-3 assay

Caspase-3 expression level was detected in MDA-MB-231 untreated and camel recombinant IFNε treated cell line using Human Caspase-3 (Casp-3) sandwich ELISA Kit (SinoGeneClon Biotech Co., Ltd) according to manufacturer’s instructions.

## Results and Discussion

### *C. dromedarius* IFNε full-length cDNA isolation and sequence analysis

By far, most information about type I IFNs has stemmed from the studies of IFNs from other species such as human, turkey, zebra fish, and bovine, but no published data is available on the Arabian camel IFNs [13, 22, 23]. In the present study, the full-length IFNε cDNA of the Arabian camel, *C. dromedarius*, was isolated by RT-PCR using gene-specific primers designed from the available expressed sequence tag (EST) camel genome project database (http://camel.kacst.edu.sa/). The PCR product corresponding to the 582 nucleotides represents the full-length IFNε cDNA (Fig 1). The PCR product was cloned into the pGEM-T-Easy vector, and the cDNA insert was sequenced using the T7 and SP6 primers. The nucleotide sequence was deposited in the GenBank database under the accession number MHO25455. Comparing the nucleotide sequence of the Arabian camel IFNε cDNA with the nucleotide sequences of other species deposited in the GenBank database using the Blastn and Blastp programs available on the National Center for Biotechnology Information (NCBI) server revealed that the putative camel IFNε gene has high statistically significant similarity scores to numerous IFNε genes from other species (Table 1). To determine the relatedness of *C. dromedarius* IFNε with known amino acid sequences available in the GenBank database, a multiple sequence alignment was conducted (Fig 2). It was observed that the percentage identity was 100% for *Camelus ferus* (GenBank accession no. XP_006179655), 95% for *Vicugna pacos* (GenBank accession no. XP_006215195), 82% for *Sus scrofa* (GenBank accession no. NP_001098780), 78% for *Bos taurus* (GenBank accession no. XP_005209958), and 75% for *Homo sapiens* (GenBank accession no. NP_795372). Moreover, the camel IFNε has high amino acid sequence homology with other type I IFNs, of which the closest paralog is IFNβ, and they share 38% identical residues [12]. A phylogenetic tree constructed (Fig 3) from the amino acid sequences of the predicted IFNε proteins deposited in the GenBank indicated that the Arabian camel IFNε took a separate evolutionary line distinct from other ungulates and mammalian species, including *H. sapiens*.

**Table 1.**
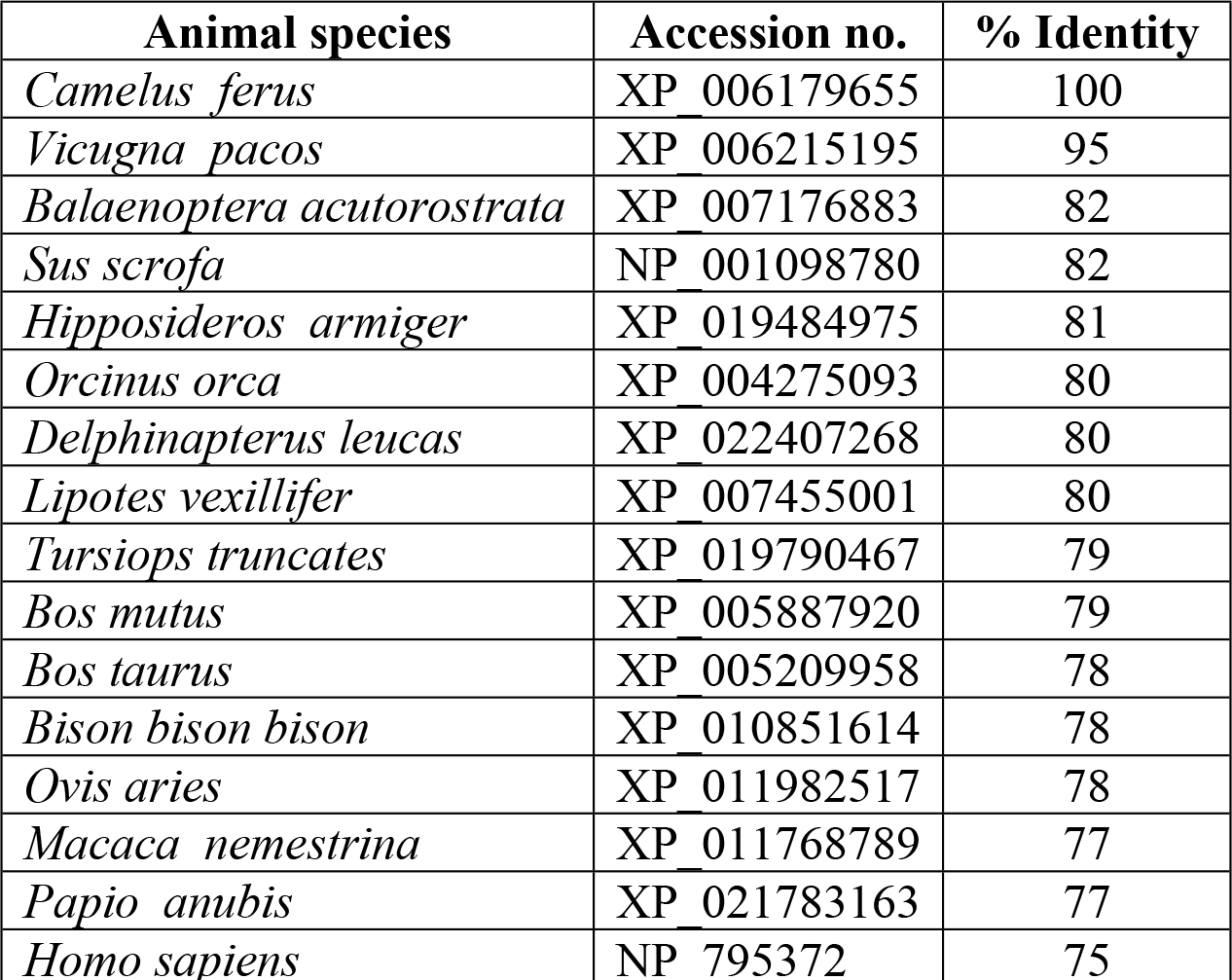
Homology of the deduced amino acids of *C. dromedarius* interferon epsilon with other species.

| Animal species | Accession no. | % Identity |
| --- | --- | --- |
| *Camelus ferus* | XP_006179655 | 100 |
| *Vicugna pacos* | XP_006215195 | 95 |
| *Balaenoptera acutorostrata* | XP_007176883 | 82 |
| *Sus scrofa* | NP_001098780 | 82 |
| *Hipposideros armiger* | XP_019484975 | 81 |
| *Orcinus orca* | XP_004275093 | 80 |
| *Delphinapterus leucas* | XP_022407268 | 80 |
| *Lipotes vexillifer* | XP_007455001 | 80 |
| *Tursiops truncatus* | XP_019790467 | 79 |
| *Bos mutus* | XP_005887920 | 79 |
| *Bos taurus* | XP_005209958 | 78 |
| *Bison bison bison* | XP_010851614 | 78 |
| *Ovis aries* | XP_011982517 | 78 |
| *Macaca nemestrina* | XP_011768789 | 77 |
| *Papio anubis* | XP_021783163 | 77 |
| *Homo sapiens* | NP_795372 | 75 |

**Fig 1.**
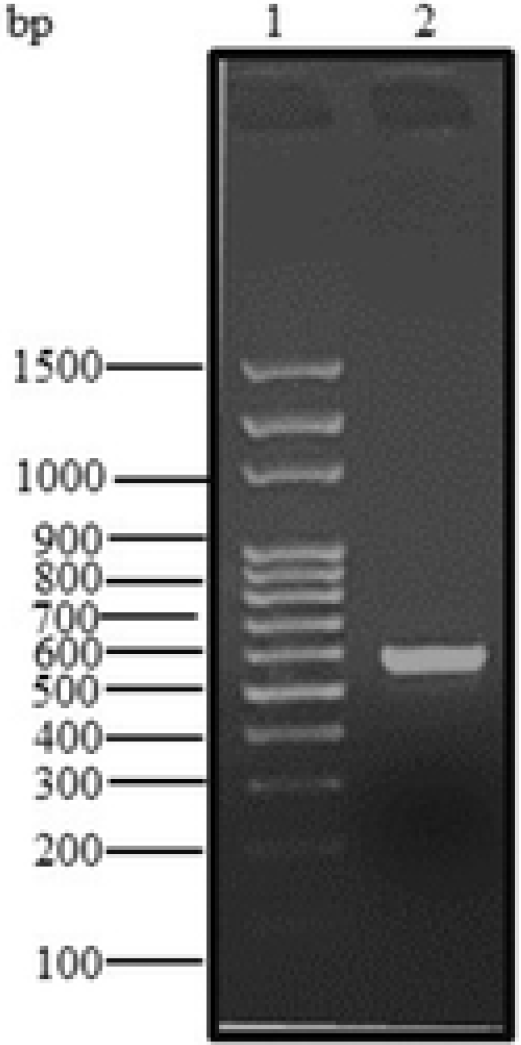
Agarose gel (1.5%) electrophoresis of PCR product for *C. dromedarius* interferon epsilon gene (Lane 2). Lane 1 represents 100 base pair DNA ladder.

**Fig 2.**
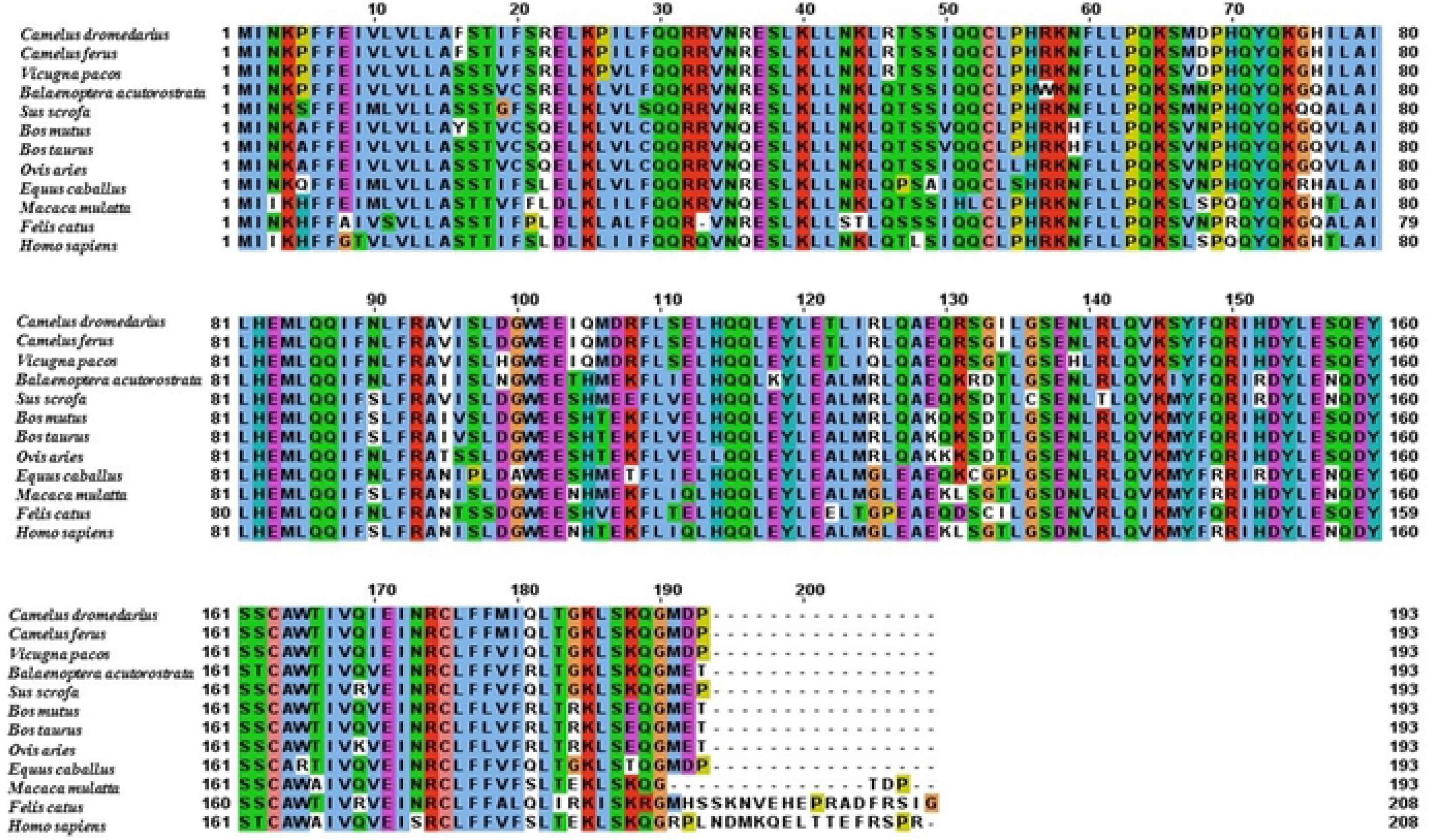
Alignment of the deduced amino acid sequence of *C. dromedrius* interferon epsilon with other species.

**Fig 3.**
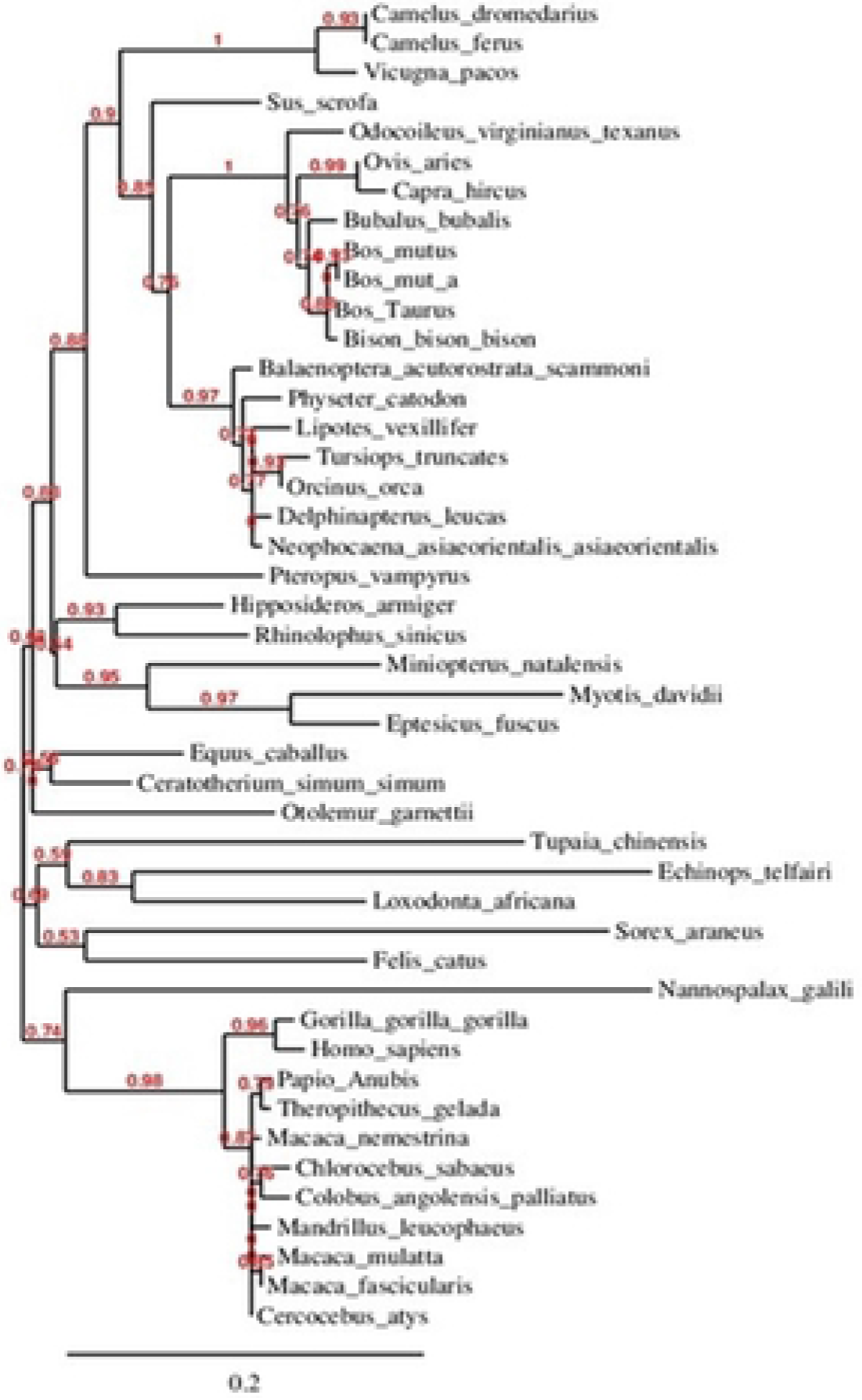
Phylogenetic relationship of *C. dromedarius* interferon epsilon and sequences from other species. Maximum likelihood tree based on complete coding sequences deposited in GenBank. Values at nodes are bootstrap ≥50%, obtained from 1000 re-samplings of the data.

### *C. dromedarius* IFNε structure annotations and predicted 3D structure

The Arabian camel IFNε primary structure and the protein motif secondary structure annotation prediction are shown in Figs 4 and 5. The nucleotides and the deduced amino acid sequence showed an open reading frame consisting of 582 nucleotides and 193 amino acid residues with a calculated molecular weight of 22.953 kDa. The isoelectric point, predicted using a computer algorithm, was found to be 9.03. From the primary structure and the multiple sequence alignment of camel IFNε with other ungulates and human, several observations merit discussion. First, the primary structure homology was greater than 75% among type I IFNs of different species. The high degree of amino acid sequence identity and conservation is presumably due to the functional constraints during evolution, although it was clear from the phylogenetic tree analysis (Fig 3) that the camel IFNε took a separate evolutionary line away from other species having type I IFNs. Second, the putative Arabian camel IFNε protein is characterized by the presence of amino acid residues Ser^38^, Glu^112^, and Ile^167^ that are highly conserved among type I INFε. Third, the Arabian camel IFNε putative protein contained three cysteine residues (Cys^53^, Cys^163^, and Cys^175^), like those found in the bovine IFNε, and two of them, probably Cys^53^ and Cys^175^, might be involved in the formation of intramolecular disulfide bonds that link the N-terminus to the end of helix F. These cysteine residues are highly conserved amongst other members of the examined type-I IFN homologs, such as human IFNλ and β and rabbit interferon-γ [38–40]. Fourth, the analysis of putative glycation sites in the camel IFNε protein (Figs 3 and 4) led to the prediction of seven potential glycation sites, although not occurring within the conserved signal for glycosylation, Asn-Xaa-Ser/Thr; these sites are 3NKPF, 35NRES, 43NKLR, 59NFLL, 90NLFR, 139NLRL, and 173NRCL. These glycation sites might act as the sites of protection against proteases-mediated hydrolysis and contributing to the process of folding, oligomerization, and stability of the protein. The identification of such sites raised the possibility that the putative camel IFNε might form a glycoprotein [41]. Fifth, the Arabian camel IFNε amino acid sequence was characterized by the presence of IFNAR-1- and IFNAR-2-binding domains. The putative IFNAR-1-binding domain is critical for receptor recognition and biological activity, and this domain was represented by the amino acid residues, F^29^, Q^30^, R^33^, R^36^, E^37^, K^40^, N^43^, and K^44^, located in the first α-helix of the camel IFNε protein (Fig 5). The IFNAR-2-binding site contained the amino acid residues, L^54^, P^55^, H^56^, R^57^, K^58^, N^59^, F^60^, L^61^, P^63^, Q^64^, K^65^, Q^71^, and Y^72^. Other conserved amino acids residues involved in the binding of different ligands and DNA are shown in Table 2.

**Fig 4.**
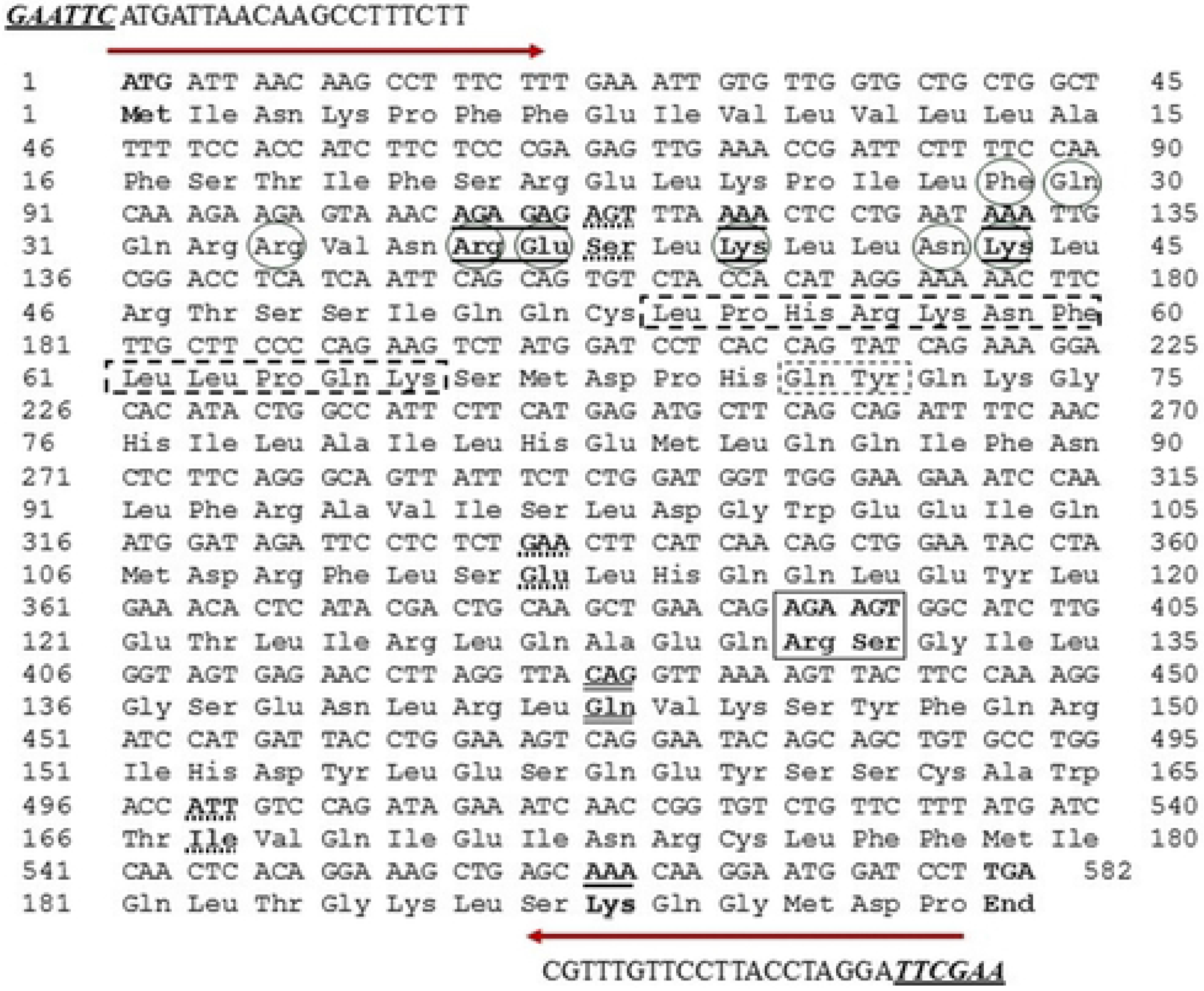
Nucleotide and deduced amino acid encoding region of *C. dromedarius* IFNε. Important amino acid residues and regions include: residues contact to N-Acetyl-2-Deoxy- are in box; residues contact to SO4 ion are in bold underline; residue contact to Zn ^+2^ are bold double underline, conserved amino acid residues in IFNε protein is in bold dashed underline, residues involved in IFNAR-1 binding are in circle and residues involved in IFNAR-2 binding are in bold dashed box. Arrows indicates the location of the forward and reverse primers with restriction enzyme sites are in bold underline italics.

**Fig 5.**
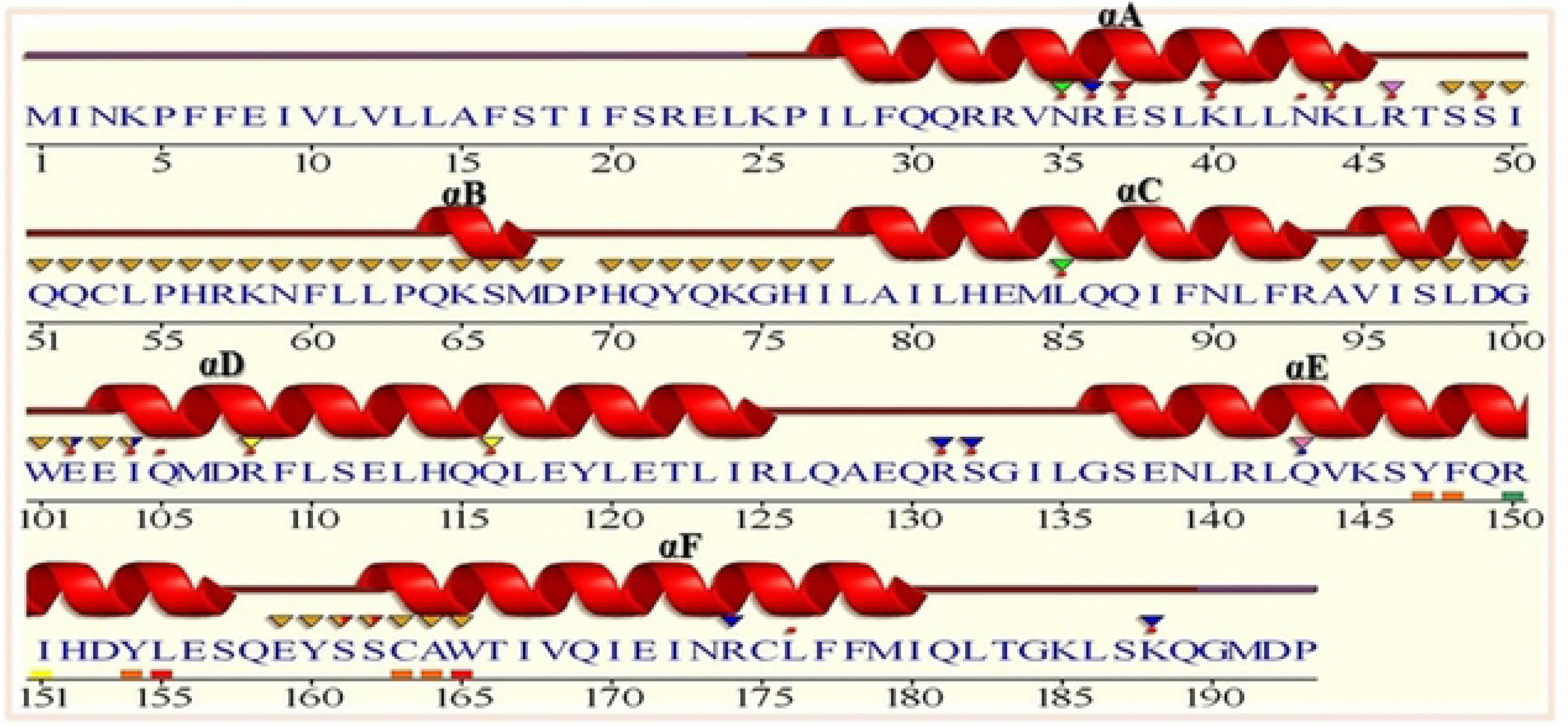
Sequence annotations for *C. dromedarius* IFNε showing the location of α-helices and residues contact to ligand and ions. Secondary structure by homology 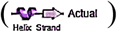 active sites residues from PDB site record 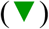; residues contacts to ligand 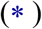 and to ions 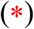.

**Table 2.** Conserved amino acid residues of *C. dromedarius* interferon ε involved in different ligands and metal ions binding.

| Annotation features | Amino acid residues |
| --- | --- |
| Contact(s) to ligands<br>- N-Acetyl-2-Deoxy-2-Amino-Galactose | Arg<sup>131</sup>, Ser<sup>132</sup> |
| Sulfate ion (SO<sub>4</sub>) | Arg<sup>36</sup>, Glu<sup>37</sup>, Lys<sup>40</sup>, Lys<sup>44</sup>, Lys<sup>188</sup> |
| - Beta-D-Glucose, 6-Deoxy-Alpha-D-Glucose | Lys<sup>44</sup>, Glu<sup>102</sup>, Ile<sup>104</sup>, Gln<sup>105</sup>, Arg<sup>108</sup> |
| - 1,2-Ethanediol | Asn<sup>35</sup>, Asn<sup>43</sup>, Arg<sup>46</sup>, Leu<sup>85</sup>, Gln<sup>116</sup>, Leu<sup>176</sup> |
| - 4-(2-Hydroxyethyl)-1-Piperazine | Arg<sup>46</sup>, Ser<sup>49</sup> |
| Contact(s) to metals<br>-Zinc ion | Gln<sup>143</sup> |
| Contact(s) to nucleic acids | Gln<sup>30</sup>, Arg<sup>33</sup>, Val<sup>34</sup>, Arg<sup>36</sup>, Glu<sup>37</sup>, Leu<sup>39</sup>, Lys<sup>40</sup>, Asp<sup>107</sup>, Ser<sup>111</sup>, Glu<sup>112</sup>, Gln<sup>115</sup>, Glu<sup>118</sup>, Tyr<sup>119</sup>, Phe<sup>177</sup>, Gln<sup>181</sup>, Gly<sup>184</sup> |

The predicted 3D structure of the Arabian camel IFNε indicated that the protein secondary structure consisted of six α-helices labelled from A to F. The composition of the predicted secondary structure revealed 61.5% α-helices, 32.6% coils, and 2.1% turns. Compared with other type Iα IFNs, the camel protein showed an extended C-terminus (Fig 6 A, B, and C). It was observed that the overall folding in the 3D structure of camel IFNε was quite similar to that of the bovine-type IFNε [10]. Moreover, the alignment template model (Fig 7 A and B) showed 36.36% similarity between the camel IFNε and *H. sapiens* type-I α2 IFN, with the preservation of the components of the secondary structures, α-helices, coils, and turns.

**Fig 6.**
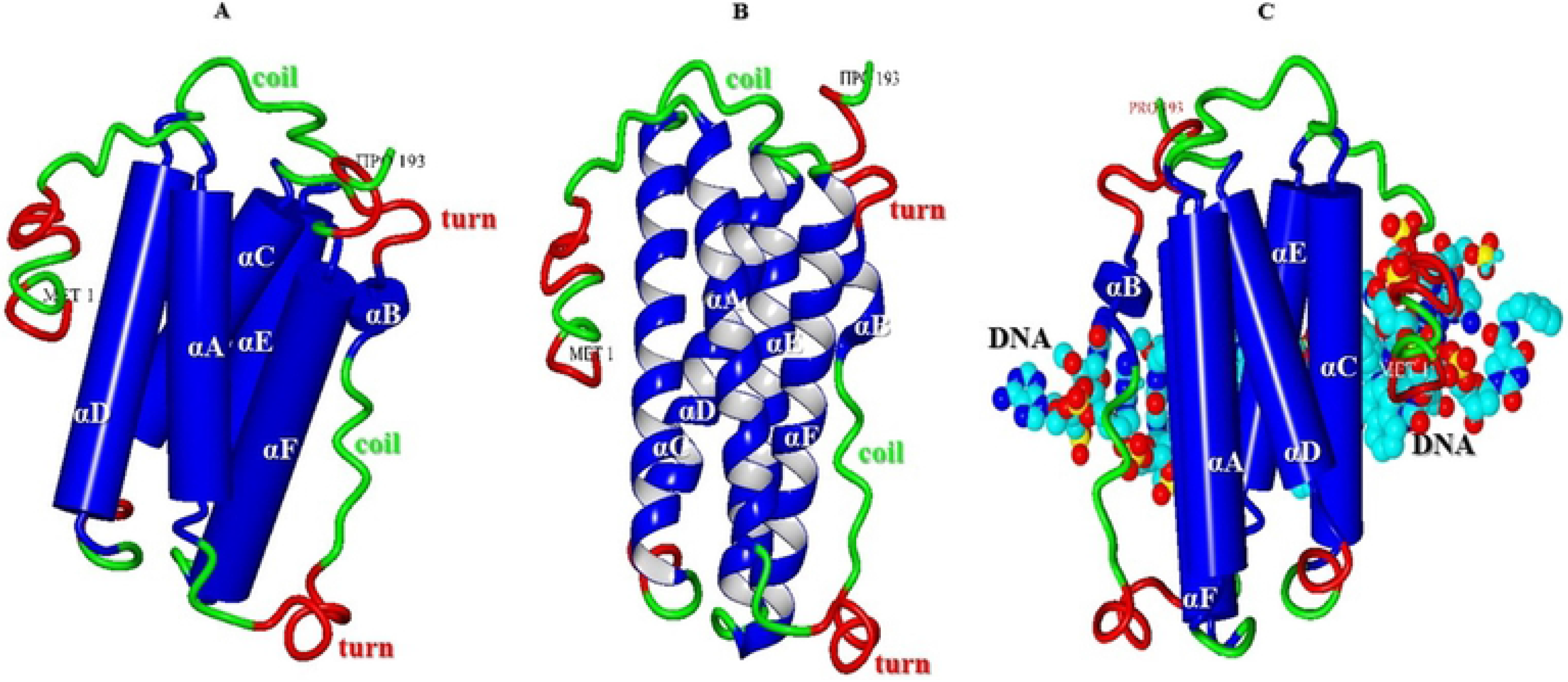
Predicted 3D structure of *C. dromedarius* IFNε protein. (A) The overall secondary structure in cartoon form, ribbon form (B) and DNA binding form (C). Components of secondary structure are α-helices (blue), coils (green) and turns (red). Alpha helices are labelled from A to F.

**Fig 7.**
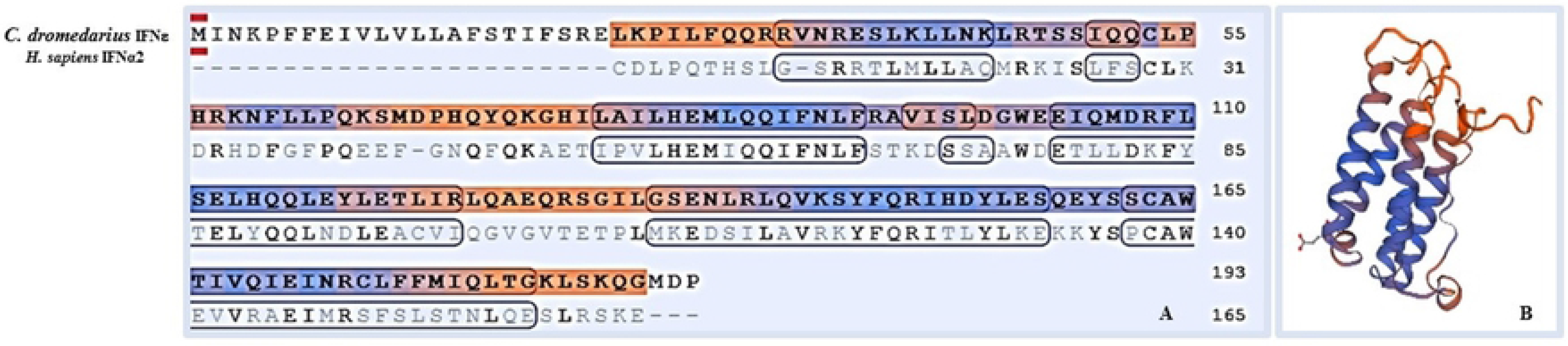
(A) Model-template alignment of amino acid residues of *C. dromedarius* IFNε and *H. sapiens* IFNα2. Components of the secondary structure are shown in blue (α-helices) and brown (coils). Identical amino acid residues are in bold black. (B) Predicted 3D structure model of *C. dromedarius* based on this model template alignment.

### Expression, solubilization, and refolding of camel IFNε protein

The Arabian camel IFNε cDNA was expressed in *E. coli* BL21(DE3) pLysS as a 6-histidine fusion protein under the control of the T7 promoter of the pET28a (+) vector. The recombinant protein was found to be overexpressed when *E. coli* cells were induced with either 1.0 mM IPTG or 2.0 g/L lactose in the fermentation medium (Fig 8 A and B). Surprisingly, the recombinant protein was found as insoluble inclusion bodies that were precipitated in the form of submicron spherical proteinaceous particles upon cell disruption by sonication and after centrifugation at 12,000 rpm for 10 min at 4 °C, leaving behind a supernatant devoid of the recombinant IFNε protein (Fig 8B). The transmission electron micrograph (Fig 9A) showed that the *E. coli* cells becomes to form dark, dense spot areas in the cytoplasm when induced to express the recombinant IFNε protein either by IPTG or lactose. The recombinant camel IFNε inclusion bodies appeared as homogeneous spherical particles of the diameter ranging from 0.5 to 1.0 μm under SEM (Fig 9 B and C). It is well documented that the expression of a foreign gene in the *E. coli* cells results in the accumulation of recombinant proteins in the form of inactive, insoluble aggregates of inclusion bodies. Thus, the biggest challenge remaining is the recovery of soluble and functional active recombinant protein from inclusion bodies; this requires standardization protocols for solubilization, re-folding, and subsequent purification [41]. Interestingly, the camel IFNε inclusion bodies are localized preferentially in the polar region of the *E. coli* cells, as well as in the mid-cell region. This polar distribution is mainly attributed to macromolecular crowding in the nucleoid region that is rich in nucleic acids and other macromolecules, which might prevent the accumulation of large protein aggregates [41]. In most cases, urea at a high concentration (4-8 M) or guanidine hydrochloride was used to solubilize and refold inclusion bodies. Our attempt to solubilize and refold the camel IFNε inclusion bodies was failed (data not shown). Thus, the alternative solubilization and refolding protocol was applied based on a strong anionic detergent SDS, which can be easily removed by precipitation with KCl. The recombinant camel IFNε inclusion bodies were collected, solubilized and refolded by the SDS/KCl method (Fig 10A, Lane 3). The solubilized and refolded inclusion bodies were subjected to nickel-affinity chromatography. The recombinant camel IFNε was bound to the affinity matrix and eluted using imidazole at a concentration of 500 mM (Fig 10 B, Peak 2). The purified protein showed a specific, unique protein band at 22.953 kDa as shown in Fig 10 C and D.

**Fig 8.**
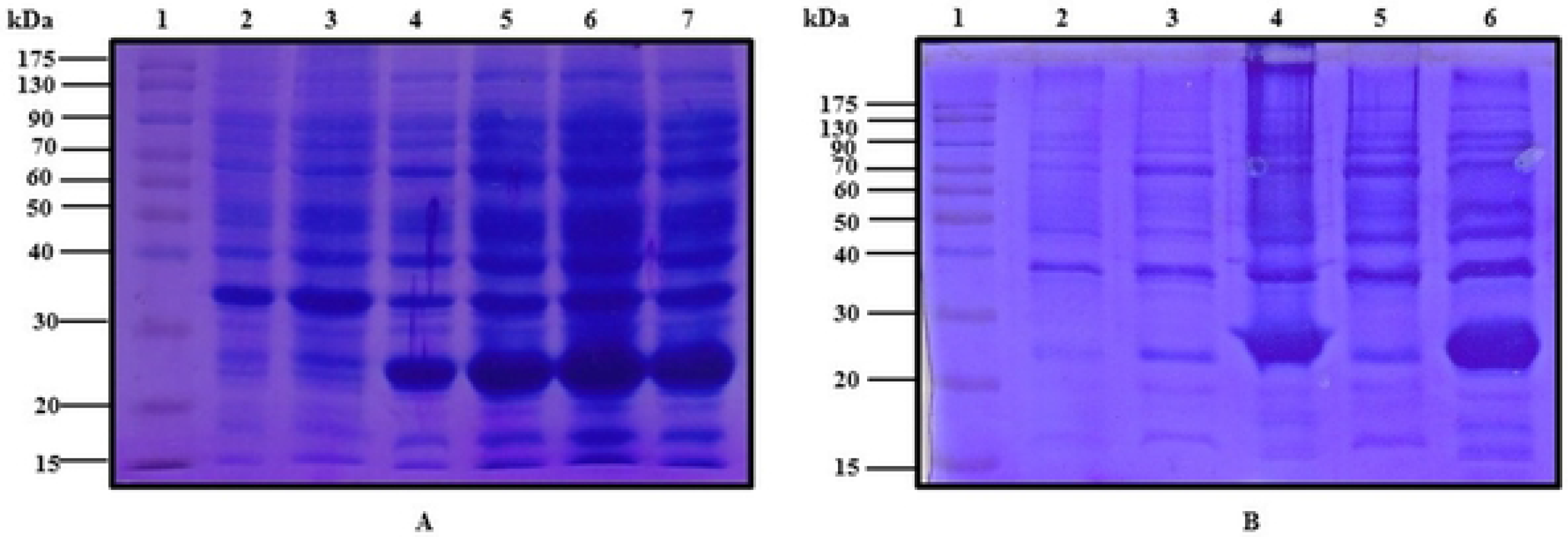
(A) SDS-PAGE (12%) for un-induced *E. coli* DE3 (BL21) pLysS pET28-a (+) harboring *C. dromedarius* IFNε cDNA (Lanes 2 and 3) and lactose induced culture (Lanes 4-7). (B) SDS-PAGE (12%) for un-induced *E. coli* DE3 (BL21) pLysS pET28-a (+) harboring *C. dromedarius* IFNε cDNA (Lane 2), IPTG induced culture supernatant (Lane 3), IPTG induced culture inclusion bodies (Lane 4), lactose induced culture supernatant (Lane 5) and lactose induced inclusion bodies (Lane 6). Lane 1 represents pre-stained protein molecular weight markers. Induction was carried out for 5 h at 1 mM IPTG and 2 g/L lactose in the fermentation medium. Arrow indicates the location of inclusion bodies.

**Fig 9.**
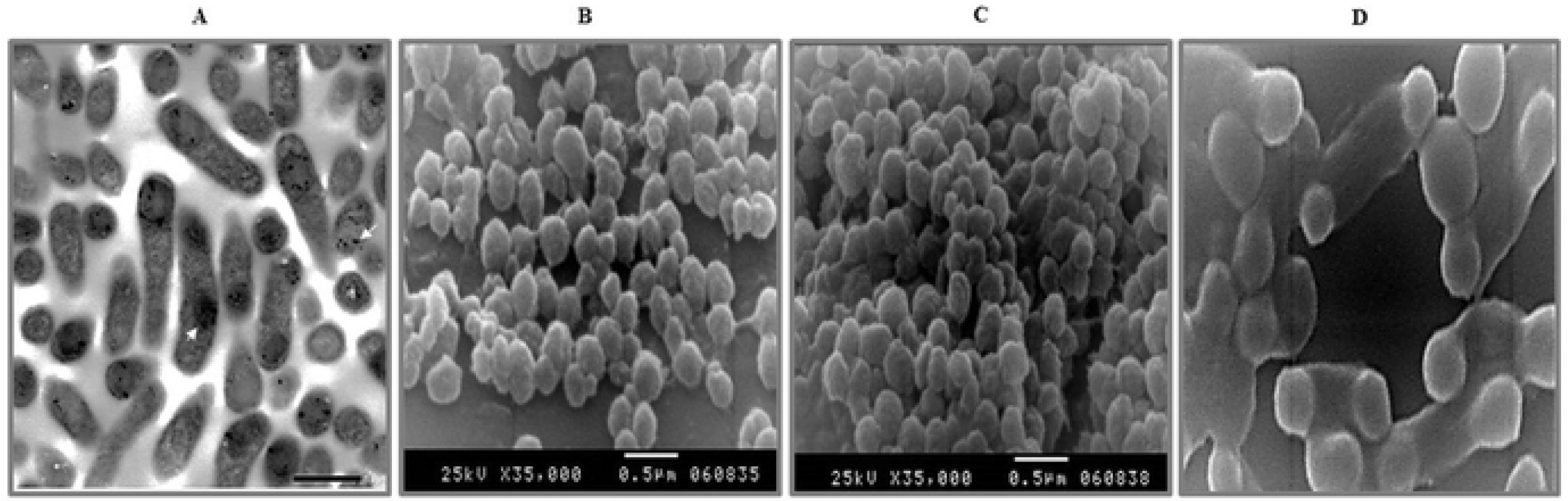
(A) Transmission electron microscope micrograph for normal *E. coli* BL21 (DE3) pLysS harboring pET28a (+) carrying *C. dromedarius* IFNε gene becomes to form inclusion bodies, dark spots when induced to overexpress the recombinant protein. Direct magnification was 10,000 x. (B), (C) and (D) Scanning electron micrograph for the inclusion body showing a spherical particle of a diameter ranging from 0.5 to 1.0 μm. Direct magnification was 35,000 x for B and C and 50,000 x for D.

**Fig 10.**
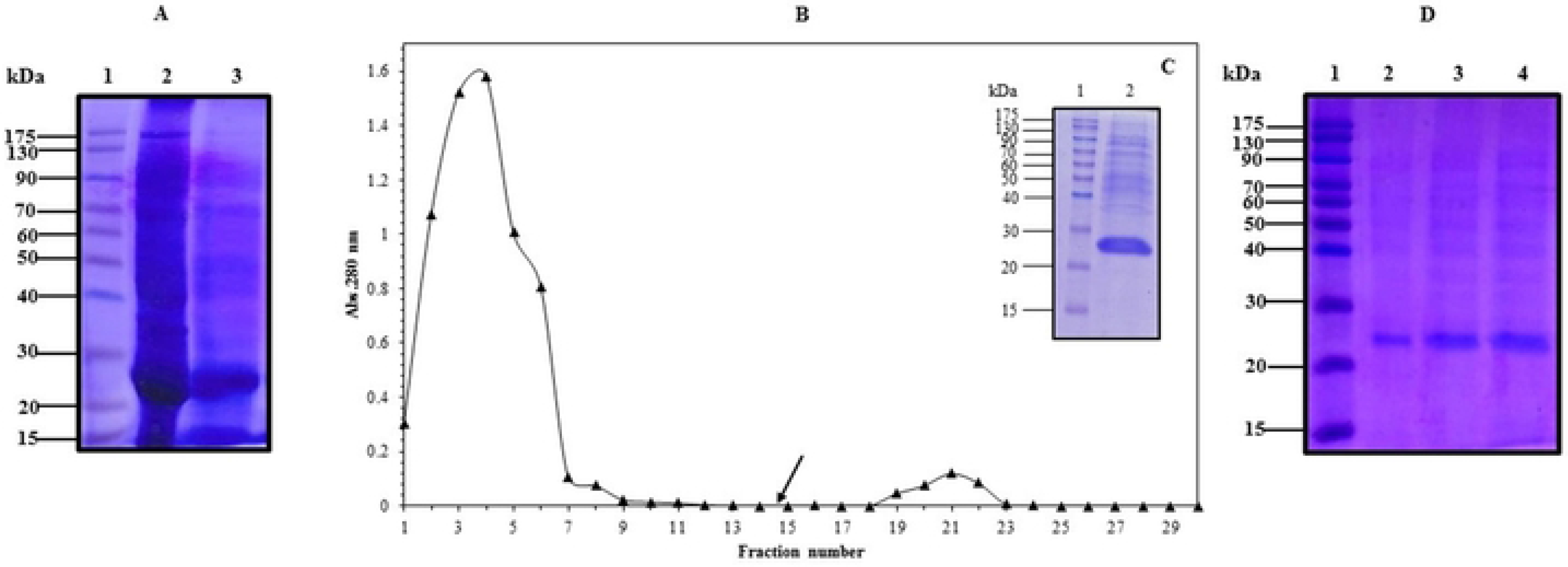
(A) SDS-PAGE of *C. dromedarius* IFNε inclusion bodies (Lane 2) and solubilized inclusion bodies (Lane 3). (B) Elution profile of *C. dromedarius* recombinant IFNε after nickel affinity chromatography. Column flow rate was adjusted to be 3 mL/5 min. Arrow indicates the fraction at which buffer was changed to contain imidazole at a concentration of 500 mM as eluent. (C) SDS-PAGE (12%) electrophoresis of nickel affinity purified refolded *C. dromedarius* IFNε, fraction # 21 (Lane 2). (D) SDS-PAGE (12%) for nickel affinity purified recombinant *C. dromedarius* IFNε (Lanes 2-4, 5-15 μg purified protein was loaded into each well). Lane 1 represents pre-stained protein molecular weight markers.

### *C. dromedarius* IFNε inhibits survival of breast cancer cells

A growing body of evidence demonstrates the antitumor effect of type I interferons [42] however, the effects of recombinant IFNε on human cancer cells have not been fully elucidated. In order to study the effects of the Arabian camel IFNε on human cancer cells, MDA-MB-231 and MCF-7 breast cancer cells were treated with different concentrations of recombinant IFNε protein. After 48 h of treatment, morphological changes were observed starting from 2.6 μM of the recombinant protein. Cells rounded up and were more easily detached. The cells exhibited shrinkage and reduction in size compared to the control cells, suggesting inhibition of cell viability (Fig 11). To investigate the effect of recombinant IFNε on cell viability, MTT assays were performed. Results demonstrate that IFNε inhibits the viability of both cell lines in a dose dependent manner. IC50 was calculated revealing concentrations of 5.65±0.2μM and 3.91±0.6 μM for MDA-MB-231 and MCF-7 cells, respectively (Fig 12). Evasion of regulated modes of cell death has been well established as a hallmark of cancer [43]. To understand the mechanism underlying IFNε-induced inhibition of cancer cell survival, MDA-MB-231 and MCF-7 cells were incubated with IFNε, and apoptosis assays were performed. Results reveal that, IFNε induces early and late apoptosis in both cell lines (Fig 13). Taken together interferon epsilon induces morphological changes and inhibits the survival of cancer cells in a dose dependent manner via the induction of apoptosis. Cancer is considered an aberrant tissue/organ comprising a hierarchical composition of heterogeneous cell populations. The tumor microenvironment and related cytokines, such as interferons, play a crucial role during tumor development and regulation of cancer cell survival and tumor progression [42]. Type I INFs, such as IFNε, signal through interferon α/β receptor (IFNAR) which is composed of two subunits, INFAR1 and IFNAR2. Studies have reported that mice with an impaired Type 1 interferon signaling (*Ifnar1−/−*) are more tumor-prone compared with wild type mice when exposed to the carcinogen methylcholanthrene [44] and mice lacking functional Type I IFN signaling have shown enhanced susceptibility of for *v-Abl*-induced leukemia/lymphoma [45]. IFNAR1-deficient tumors are rejected when transplanted into wild type mice, however, tumors grow when transplanted in *Ifnar1−/−* mice, demonstrating the role of type I IFNs in carcinogenesis and tumor progression [44]. IFN-α/β has direct effects in tumor cells, inducing growth arrest and apoptosis via activating the JAK-STAT pathway and the expression of genes whose promoters contain the IFN-stimulated response element, such as the apoptosis mediators FAS and TRAIL [46, 42]. The effects of type I IFNs on cancer cells vary depending on the type of tumor, and not all tumor cells are susceptible to the apoptotic effects of IFNs. Similar to orthologs in other species, recombinant canine IFNε has shown to be capable of activating the JAK-STAT pathway and inhibiting the proliferation of canine cell lines [47]. To complement what has been investigated in the study, the expression level of Caspase-3 was determined to evaluate the cytotoxicity strength and the effectiveness of the potential camel IFNε protein. Caspase-3 expression has been directly correlated with apoptosis because of its location in the protease cascade pathway as it is activated by diverse death-inducing signal such as chemotherapeutic agents [48, 49]. Our results showed that caspase-3 expression level was increased in MDA treated cells and the fold of induction was found to be 168.03% and 157.8 % at a protein concentration of 3 and 6 μM, respectively compared to untreated control cells (Fig 14). This finding has important clinical implication and in conjunction with other studies suggest that IFNε can be considered as a chemotherapeutic agent that may help in improving the response of adjuvant therapy for breast cancer.

**Fig 11.**
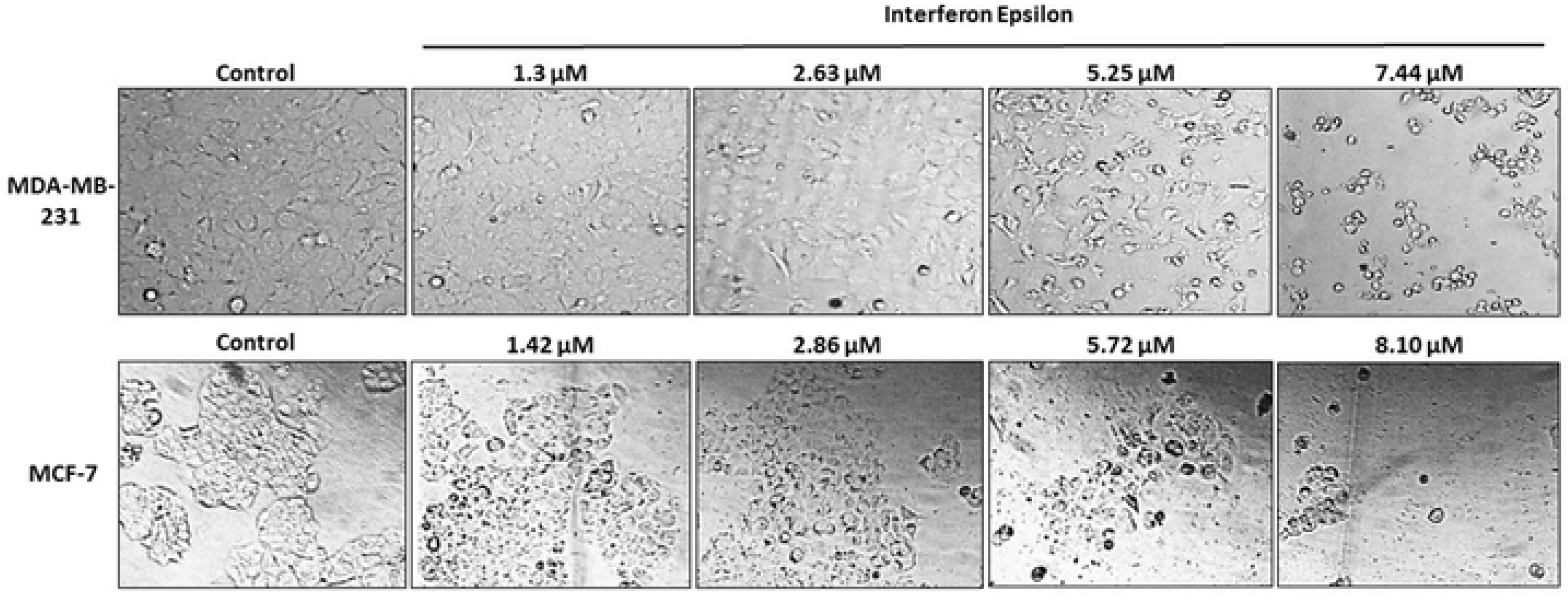
Recombinant Arabian camel IFNε alters the morphology of breast cancer cell lines MDA-MB-231 (upper) and MCF-7 (lower).

**Fig 12.**
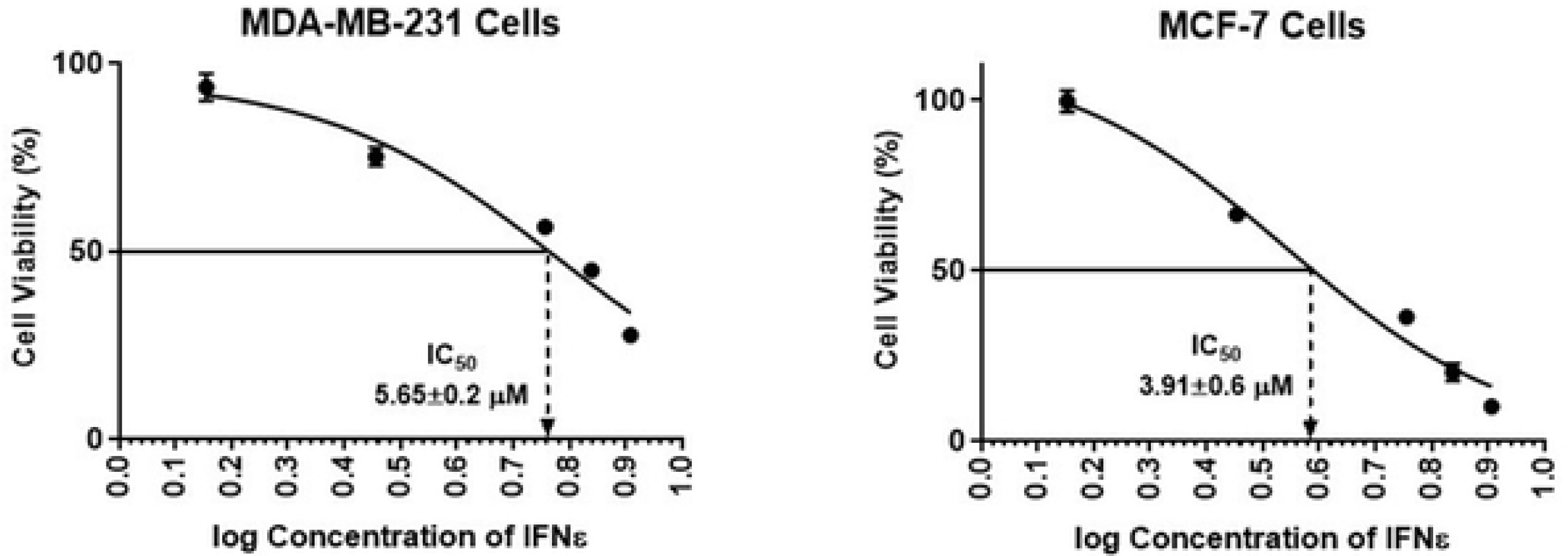
Interferon epsilon inhibits the survival of breast cancer cells.

**Fig 13.**
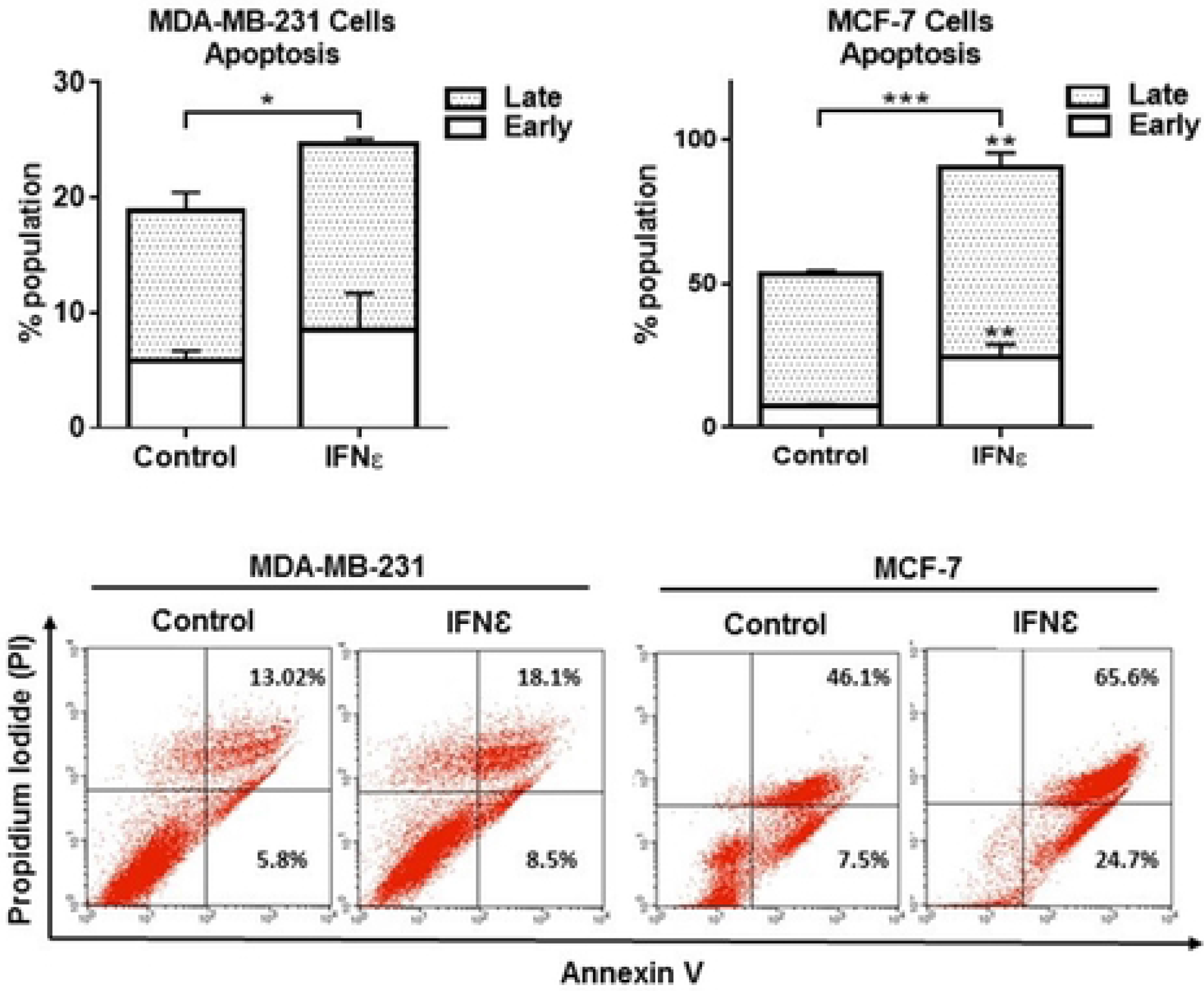
Interferon epsilon induces apoptosis in breast cancer cells. Cells were treated with 5 μM IFNε protein for 48 h. Apoptosis assay was performed, and the percentage cell viability was calculated (*p<0.5, **p<0.1 and ***p<0.01).

**Fig 14.**
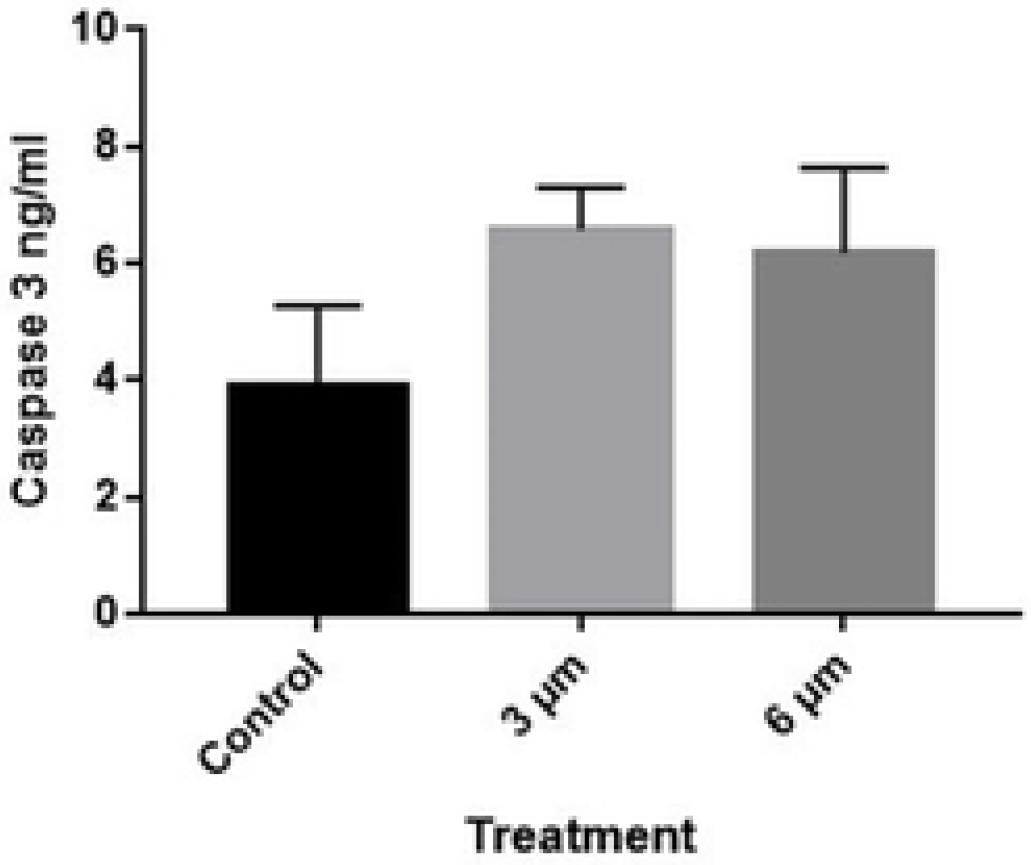
Expression of caspase-3 in MDA-MB-231 cell line untreated and recombinant IFNε treated cells at a concentration of 3 and 6 μM.

In conclusion, we present here cloning, expression, refolding, and characterization of a novel gene encoding the Arabian camel IFNε. Moreover, this study does underpin the Arabian camel recombinant IFNε as a possible novel and effective agent for the treatment of cancer.

